## Supplementary Table 1 for "Cell-selective proteomics reveal novel effectors secreted by an obligate intracellular bacterial pathogen"

| Strain or plasmid | Genotype or feature | Reference or source |
|---|---|---|
| ***R. parkeri* strains** | | |
| *R. parkeri* str. Portsmouth | Parental *R. parkeri* strain | Chris Paddock |
| WT | pRAM18dSGA[MCS] | This study |
| MetRS* | pRL0128 | This study |
| GSK-BFP | pRL0284 | Sanderlin *et al*. (2022) |
| GSK-RARP-2 | pRL0285 | Sanderlin *et al*. (2022) |
| GSK-SrfA | pRL0368 | This study |
| GSK-SrfB | pRL0369 | This study |
| GSK-SrfC | pRL0370 | This study |
| GSK-SrfD | pRL0371 | This study |
| GSK-SrfE | pRL0372 | This study |
| GSK-SrfF | pRL0373 | This study |
| GSK-SrfG | pRL0374 | This study |
| **Plasmids** | | |
| pRAM18dSGA[MCS] | *Rickettsia* shuttle vector | Ulrike Munderloh |
| pRL0128 | MetRS* | This study |
| pRL0284 | GSK-tagged TagBFP | Sanderlin *et al*. (2022) |
| pRL0285 | GSK-tagged RARP-2 | Sanderlin *et al*. (2022) |
| pRL0368 | GSK-tagged SrfA | This study |
| pRL0369 | GSK-tagged SrfB | This study |
| pRL0370 | GSK-tagged SrfC | This study |
| pRL0371 | GSK-tagged SrfD | This study |
| pRL0372 | GSK-tagged SrfE | This study |
| pRL0373 | GSK-tagged SrfF | This study |
| pRL0374 | GSK-tagged SrfG | This study |
| pRL0375 | GST-tagged SrfC | This study |
| pRL0376 | GST-tagged SrfD(Δ766-957) | This study |
| pRL0377 | GST-tagged SrfF | This study |
| pRL0378 | 6xHis-SUMO-TwinStrep-tagged SrfD(Δ766-957) | This study |
| pRL0379 | 6xHis-SUMO-TwinStrep-tagged SrfF | This study |
| pRL0381 | empty vector control | This study |
| pRL0382 | 3xFLAG-tagged SrfA | This study |
| pRL0383 | 3xFLAG-tagged SrfB | This study |
| pRL0384 | 3xFLAG-tagged SrfC | This study |
| pRL0385 | 3xFLAG-tagged SrfD | This study |
| pRL0386 | 3xFLAG-tagged SrfE | This study |
| pRL0387 | 3xFLAG-tagged SrfF | This study |
| pRL0388 | 3xFLAG-tagged SrfG | This study |
| FCW2IB-BiP-mNeonGreen-KDEL | BiP(1-18)-mNeonGreen-KDEL | Acevedo-Sánchez *et al*. (2023) |
| pRL0389 | *Gaussia*-Dura luciferase | This study |
| pRL0390 | 3xFLAG-tagged SrfDΔPPR1 | This study |
| pRL0391 | 3xFLAG-tagged SrfDΔCC1 | This study |
| pRL0392 | 3xFLAG-tagged SrfDΔPPR2 | This study |
| pRL0393 | 3xFLAG-tagged SrfDΔTM | This study |
| pRL0394 | 3xFLAG-tagged SrfDΔCC2 | This study |

**Supplementary Table 1. Strains and plasmids used in this study.**
